# Decomposition of transcriptional responses provides insights into differential antibiotic susceptibility

**DOI:** 10.1101/2020.05.04.077271

**Authors:** Anand Sastry, Nick Dillon, Saugat Poudel, Ying Hefner, Sibei Xu, Richard Szubin, Adam Feist, Victor Nizet, Bernhard Palsson

**Affiliations:** Department of Bioengineering, University of California San Diego, La Jolla, CA, 92093, USA; Department of Pediatrics, University of California, San Diego, La Jolla, CA, USA; Skaggs School of Pharmacy and Pharmaceutical Sciences, University of California, San Diego, La Jolla, CA, USA; Novo Nordisk Foundation Center for Biosustainability, Technical University of Denmark, Lyngby, 2800, Denmark

## Abstract

Responses of bacteria to antibiotic treatments depend on their environments. Differences between *in vitro* testing conditions and the physiological environments inside patients have resulted in poor antibiotic susceptibility predictions, contributing to treatment failures in the clinic. Here, we investigate how media composition affects antibiotic susceptibility in the laboratory strain *E. coli* K-12 MG1655, and contextualize these changes through machine learning of transcriptomics data. We show that complex transcriptional changes induced by different media or antibiotic treatment can be traced back to a few key regulators. Integration of results from machine learning with biochemical knowledge reveals fundamental shifts in respiration and iron availability that may explain media-dependent differential susceptibility to antibiotics. The data generation and analytical workflow used here can interrogate the regulatory state of a pathogen under any condition, and can be extended to additional strains and organisms for which data is available.

## Introduction

Antibiotic activities are highly dependent on environmental conditions. Factors such as metabolites, buffering systems, immune components, and the presence of other antibiotics can drastically alter their activities against bacteria, as measured by changes in their minimum inhibitory concentrations (MIC) (Dorschner et al., 2006; Ersoy et al., 2017; Farha et al., 2018; Kudrin et al., 2017; Meylan et al., 2018). Despite these dependencies, standardized antibiotic susceptibility testing (AST) is performed in bacteriological media such as Cation-Adjusted Mueller Hinton Broth (CA-MHB), which poorly recapitulates the *in vivo* conditions of the host (Ersoy et al., 2017). AST in tissue culture media, such as RPMI 1640 or DMEM, more closely mimics physiological conditions and may be a better predictor of *in vivo* efficacy (Ersoy et al., 2017; Lin et al., 2015). Using transcriptomic data, we sought to understand the drivers of differential antibiotic susceptibility in these media conditions.

Prior studies have demonstrated that systems analysis of omics measurements can elucidate global metabolic responses to antibiotic treatment (Yang et al., 2019; Zampieri et al., 2017). Previously, we published a high-quality transcriptomic compendium for *Escherichia coli*, named PRECISE, containing 278 expression profiles (Sastry et al., 2019). Using independent component analysis (ICA), a machine learning method developed to separate mixed signals, we decomposed PRECISE into 92 independently modulated groups of genes, or i-modulons. Since most i-modulons exhibited high overlap with previously reported regulons, we named the i-modulons after their common regulator(s). ICA simultaneously computes activities for each i-modulon across every condition in the expression compendium, which predictably reflect the activity state of the enriched regulators.

I-modulon activities have been used to infer the cellular state beyond the transcriptome. In one case, adaptive laboratory evolution of *E. coli* in excess iron conditions resulted in multiple mutations in the oxidative stress response regulator OxyR (Anand et al., 2019). In each of these strains the SOS-response regulator LexA was no longer activated by hydrogen peroxide treatment, indicating that the mutations reduced DNA damage from reactive oxygen species through constitutive activation of OxyR. I-modulons have also been successfully extracted for additional species, disentangling the transcriptional trajectory of a *Bacillus subtilis* sporulation time-course(Rychel et al., 2020), and identifying media-specific transcriptional responses in *Staphylococcus aureus* (Poudel et al., 2020).

In the present study, we demonstrate the utility of an ICA-based workflow in decomposing the transcriptomic responses to antibiotics in the *E. coli* K-12 laboratory strain MG1655. First, we show that *E. coli* exhibits a differing responses to antibiotics across three media compositions: (1) glucose M9 minimal media, a standard media for scientific discovery; (2) cation adjusted mueller hinton broth, the standard rich media for antibiotic susceptibility testing, and (3) RPMI 1640 with 10% LB, a rich media that mimics physiological conditions. We then measured the expression profiles of *E. coli* on the three media and identified key transcriptomic differences in metabolism and stress responses using ICA. Finally, we examined the expression profiles of *E. coli* after treatment with subinhibitory concentrations of antibiotics and observed that subinhibitory antibiotic treatment in rich media resulted in decelerated respiration. Overall, this work provides a roadmap for interrogating the effects of antibiotic treatment in different environmental conditions.

## Results

### Antibiotic activities vary across media compositions

We measured the minimum inhibitory concentrations (MICs) of three antibiotics, ranging in drug class and mechanism of action, on glucose M9 minimal media (M9), RPMI 1640 + 10% LB (R10LB) and cation-adjusted Mueller Hinton Broth (CA-MHB) (Table 1). We then selected four antibiotics to further investigate. Ceftriaxone (CEF) and trimethoprim-sulfamethoxazole (T/S) had identical MICs in the rich media (CA-MHB and R10LB) and a lower MIC in M9. However, *E. coli* was most susceptible to ciprofloxacin (CIP) in R10LB least susceptible in CA-MHB, and meropenem (MER) exhibited an opposing susceptibility trend.

**Table 1:**
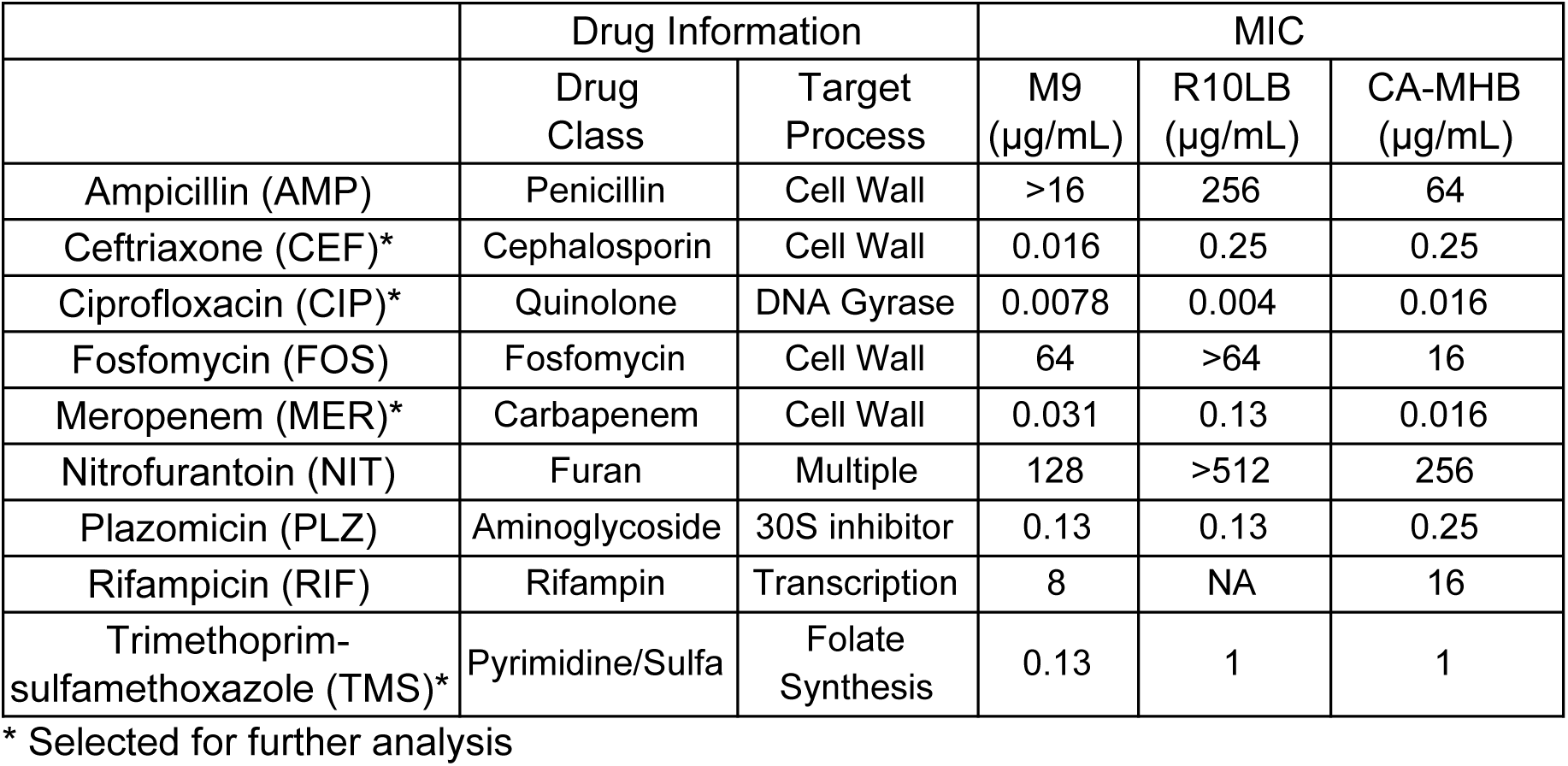
MIC of nine antibiotics on three media

|  | Drug Information |  | MIC |  |  |
| --- | --- | --- | --- | --- | --- |
| | Drug Class | Target Process | M9 ($\mu\text{g/mL}$) | R10LB ($\mu\text{g/mL}$) | CA-MHB ($\mu\text{g/mL}$) |
| Ampicillin (AMP) | Penicillin | Cell Wall | >16 | 256 | 64 |
| Ceftriaxone (CEF)* | Cephalosporin | Cell Wall | 0.016 | 0.25 | 0.25 |
| Ciprofloxacin (CIP)* | Quinolone | DNA Gyrase | 0.0078 | 0.004 | 0.016 |
| Fosfomycin (FOS) | Fosfomycin | Cell Wall | 64 | >64 | 16 |
| Meropenem (MER)* | Carbapenem | Cell Wall | 0.031 | 0.13 | 0.016 |
| Nitrofurantoin (NIT) | Furan | Multiple | 128 | >512 | 256 |
| Plazomicin (PLZ) | Aminoglycoside | 30S inhibitor | 0.13 | 0.13 | 0.25 |
| Rifampicin (RIF) | Rifampin | Transcription | 8 | NA | 16 |
| Trimethoprim-sulfamethoxazole (TMS)* | Pyrimidine/Sulfa | Folate Synthesis | 0.13 | 1 | 1 |

\* Selected for further analysis

MICs were measured in 96-well plates using an inoculum density of 1*10^5^ CFU/mL, as recommended by the Clinical and Laboratory Standards Institute (CLSI). However, we required a significantly higher cell density (~2.5*10^7^ CFU/mL) to collect enough mRNA for sequencing. Since antibiotic activity can be diluted at higher cell densities, we determined the minimum bactericidal concentration (MBC) in each media type for the targeted antibiotics at the same culture density that was used for the mRNA isolations (Figure 1a). In all conditions, the MBC was higher than the selected MICs. We therefore measured the change in MIC at RNA-seq densities, and found that the MICs were increased up to 8-fold at higher densities (Figure 1b).

**Figure 1:**
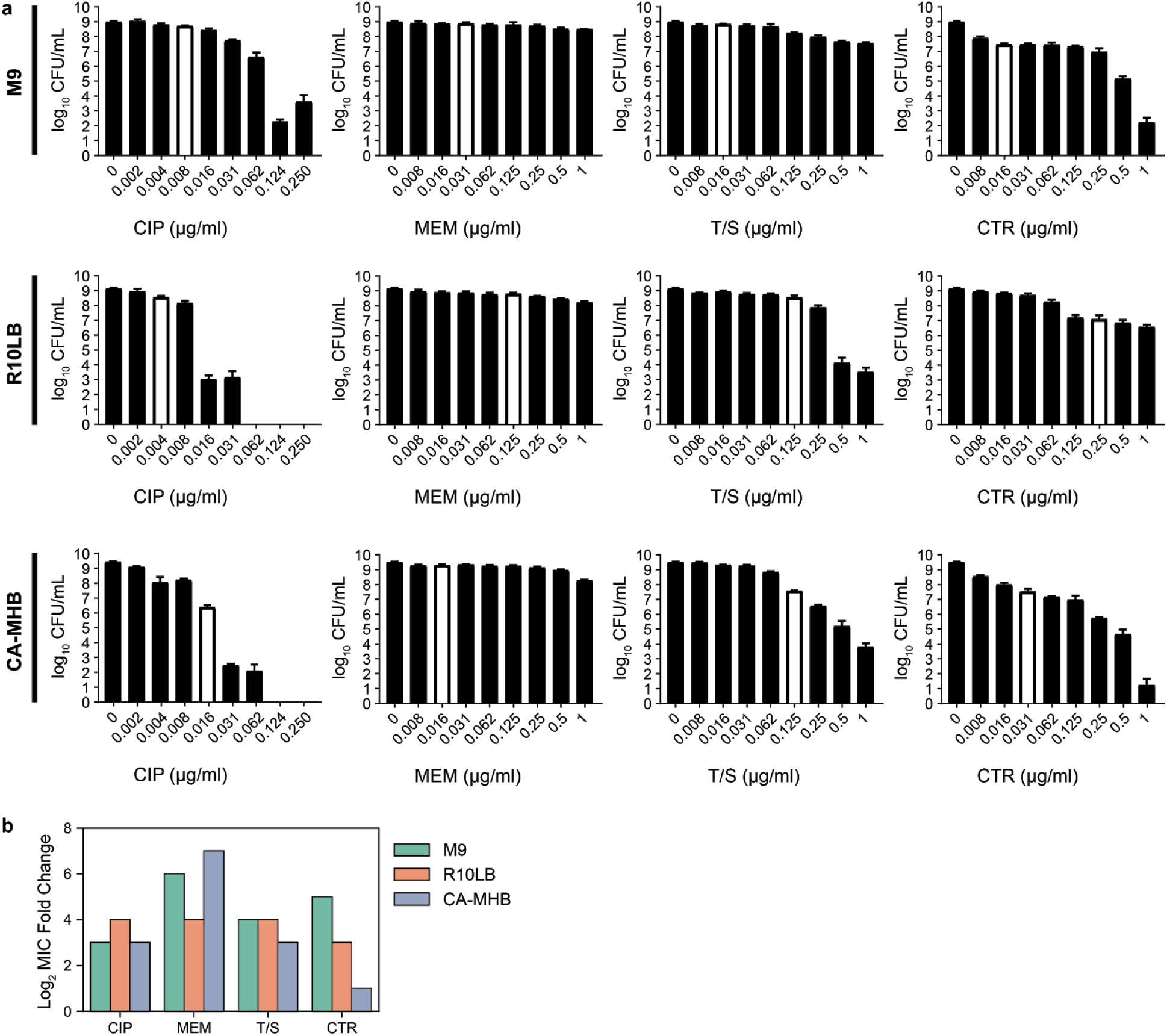
(a) MBC measurements of four antibiotics across three media compositions at gene expression conditions. White bars designate the concentration of antibiotics used in RNA-seq cultures (i.e. MIC in relevant media) (b) Changes in MIC between low (~10^5^ CFU/mL) and high (~10^7^ CFU/mL) cell densities.

### A few mechanisms can explain large transcriptomic perturbations

We first performed RNA-seq on the untreated strains grown in M9, R10LB, and CA-MHB to understand how *E. coli* responds to different media without antibiotic treatment. Pairwise differential expression analysis of the three growth conditions identified up to 1145 differentially expressed genes (DEGs) between growth conditions, comprising one quarter of all genes in *E. coli* (Figure 2a). The DEGs spanned every Cluster of Orthologous Gene (COG) category, with no known function for 21% of all DEGs. Rich media (R10LB and CA-MHB) induced similar transcriptional responses, as only 556 genes were differentially expressed between the two media.

**Figure 2:**
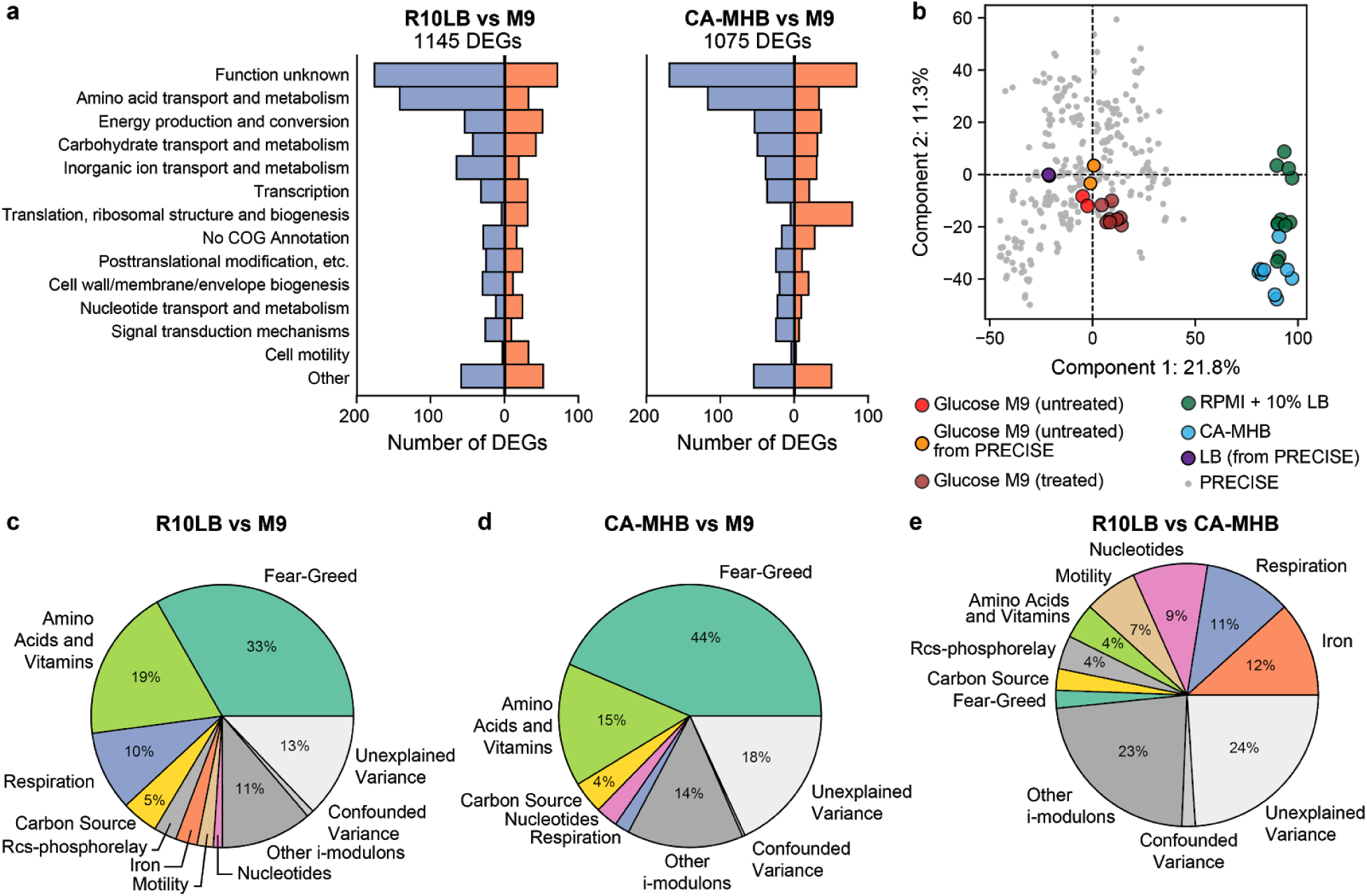
Global comparative analysis of the *E. coli* transcriptome across three media compositions. (a) Differentially expressed genes across COG categories between M9 and the two rich media (RPMI and CA-MHB). (b) Top two principal component loadings for the PRECISE dataset, comprising 278 RNA-seq profiles (Sastry et al., 2019), combined with the data generated in this study. (c-f) Proportion of expression deviation explained by 8 groups of i-modulons. I-modulons in each group are shown in Figure 3 and Figure S2c,d. Percentages above 3% are shown.

The newly-generated RNA-seq data was compared to a previously published high-quality compendium containing 278 *E. coli* expression profiles, named PRECISE, to contextualize the transcriptional response to the three media types (Sastry et al., 2019). Principal component analysis (PCA) of the combined dataset showed that CA-MHB and R10LB produced responses that were significantly different than the previously probed conditions in PRECISE, including growth on LB media without RPMI (Figure 2b). On the other hand, the expression levels on M9 minimal media were highly similar to expression levels under identical conditions produced five years ago (R^2^ = 0.85), highlighting the internal consistency of this dataset. This older expression profile grown on glucose minimal media serves as the reference condition, to which all i-modulon activities are compared.

We previously showed that independent component analysis (ICA) can be applied to large RNA-seq datasets to extract independently modulated sets of genes, termed i-modulons, and their condition-specific activities (Sastry et al., 2019). I-modulons exhibit high overlap with classically-defined regulons, which allows us to directly associate i-modulons to one or more known transcriptional regulators. Unlike many module detection methods (Saelens et al., 2018), ICA also computes i-modulon activities across all expression profiles, which represent the activity level of the linked regulator under each specific condition.

We subsequently combined the 278 datasets in PRECISE combined with 30 new RNA-seq datasets to form a compendium of 308 profiles (Supplementary Data). Application of ICA identified 98 i-modulons, of which 88 were nearly identical to i-modulons extracted from the original PRECISE dataset of 278 profiles (Sastry et al., 2019) (Figure S1). On average, i-modulons could explain 81% of the difference in expression between each pair of media conditions (Figure 2c-e). Borrowing from multiple linear regression, we also calculated how much of the expression deviation could be directly explained by each i-modulon (See Methods).

### I-modulons elucidate transcriptional markers of ‘greedy’ growth in rich media

The major source of expression variation between either rich media (RPMI and CA-MHB) and minimal media was explained by the ‘fear-greed’ tradeoff (Sastry et al., 2019) (Figure 3a). This trade-off is between fast growth (i.e. ‘greed’) and high stress response readiness (i.e. ‘fear’). In rich media, cells are significantly less stressed by their environment, leading to a strong decrease in the activity level of stress-related i-modulons, such as RpoS and the GadEWX acid-response system (Seo et al., 2015). In turn, this frees cellular resources that can be allocated to promote growth, as indicated by the increases in activity levels of i-modulons encoding genes related to translation and the central dogma of molecular biology. These two processes are molecularly linked through ppGpp, the stress alarmone, which binds to RNA polymerase to wobble the transcriptome (Sanchez-Vazquez et al., 2019). The i-modulon activities under rich media are similar to the i-modulon activities of an RpoB mutant acquired during adaptive laboratory evolution on glucose minimal media (Utrilla et al., 2016), which is hypothesized to affect the ppGpp binding site on RNA polymerase.

**Figure 3:**
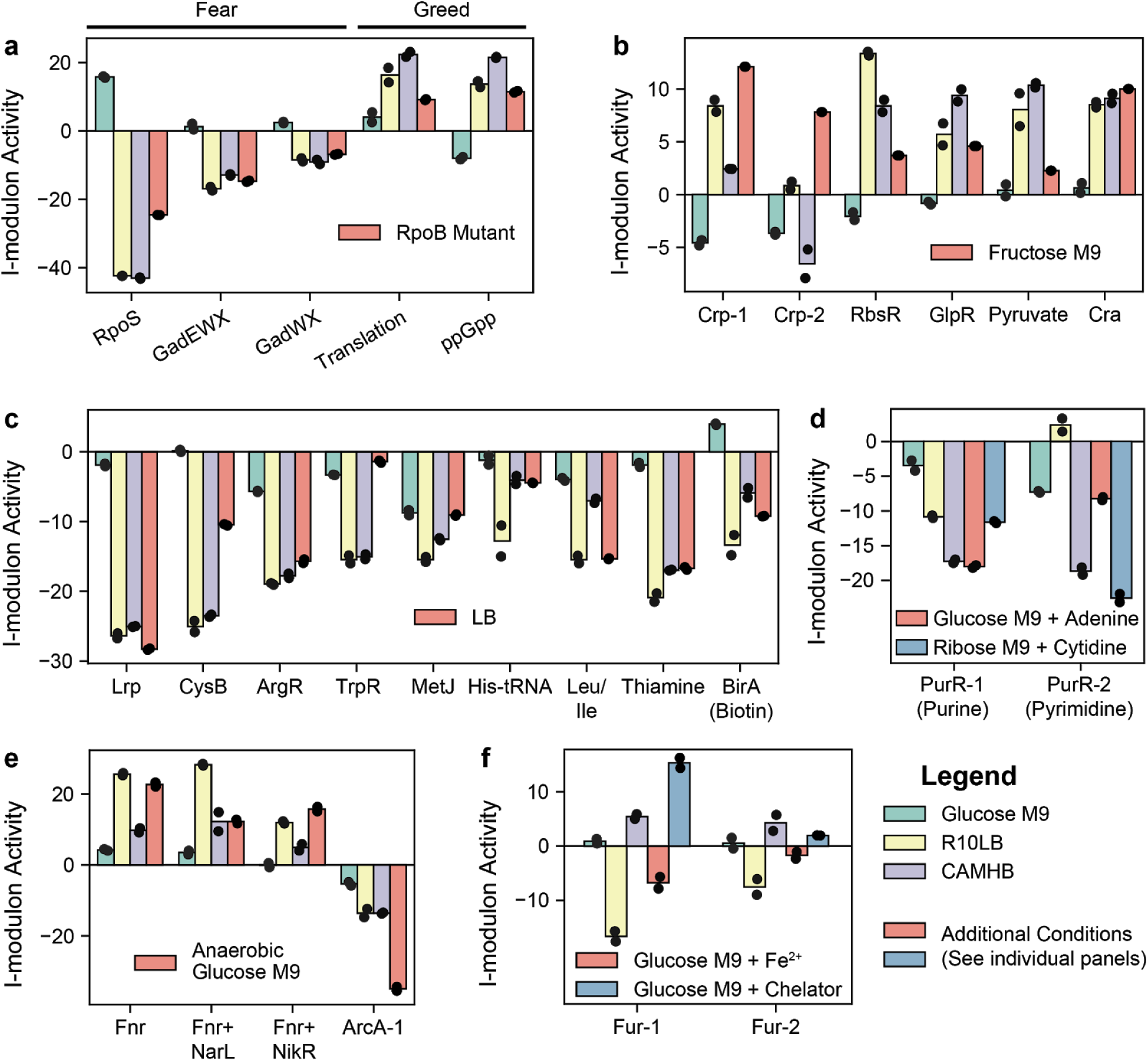
Mechanisms underlying the complex transcriptional response to different media. Bar charts show the i-modulon activities for (a) fear-greed i-modulons; (b) Carbon source catabolism i-modulons; (c) amino acid and vitamin B i-modulons; (d) Nucleotide biosynthesis i-modulons; (e) Respiration i-modulons; (f) Iron-related i-modulons. Individual measurements for independent biological replicates are plotted on top of bars.

R10LB contains glucose in its formulation, but displays surprisingly high transcriptomic similarities to glucose-starved media (Figure 3b). The cAMP receptor protein, CRP, is activated when glucose is absent from the media, recruiting RNA polymerase to genes encoding catabolic pathways for alternative carbon sources (Kolb et al., 1993). Although glucose uptake was not previously detected in CA-MHB (Poudel et al., 2020), some of its CRP-related i-modulon activities are closer to M9 media than R10LB.

Biosynthetic pathways for vitamins and amino acids contributed to 19% and 15% of the expression deviation from M9 for R10LB and CA-MHB, respectively (Figure 3c). These responses are consistent with the defined formulation of RPMI, which contains all amino acids and multiple B-vitamins, and shows that the undefined media CA-MHB and LB have a similar nutritional content to R10LB. The main deviations appear to be in histidine and branched-chain amino acid biosynthesis, which are all regulated by tRNA-mediated transcriptional attenuation. The two media also deviate with respect to nucleotides as pyrimidine biosynthesis is upregulated in R10LB, indicating that cells are starved for pyrimidines (Figure 3d). Although nucleotides are not in the RPMI formulation, the cells downregulated the *de novo* purine biosynthesis pathway.

The next largest contributor to expression deviation in R10LB included four i-modulons related to anaerobic respiration (Figure 3e). Three i-modulons were enriched with the transcription factor Fnr; one i-modulon was uniquely enriched with the Fnr regulon, one i-modulon was enriched with genes co-regulated by Fnr and the nitrate-responsive regulator NarL and the final i-modulon contained genes co-regulated by Fnr, NarL, and the nickel regulator NikR. The fourth i-modulon in this group was related to ArcAB, a two-component system that senses the redox state of quinone electron carriers to regulate the TCA cycle, among other genes (Federowicz et al., 2014; Georgellis et al., 2001). The ArcA i-modulon activity was partially reduced in both R10LB and CA-MHB.

Another major difference between the two media is the presence of free iron, as indicated by the differential activation of two Fur-enriched i-modulons (Figure 3f). Fur binds to free iron (II) and represses genes related to iron siderophore synthesis and transport (Seo et al., 2014). The Fur i-modulons are not correlated with Fur expression, and are likely driven by intracellular iron availability (Figure S2a,b). From the i-modulon activities, cells growing in glucose M9 media appear to contain more free iron than CA-MHB, but significantly less than R10LB.

A few additional i-modulons were differentially activated across the multiple media conditions, and coincidently discriminate the biological processes that are perturbed between the old and new glucose M9 expression profiles. The new glucose M9 and CA-MHB expression profiles displayed an activation of three i-modulons related to the Rcs-phosphorelay (Figure S2c), whereas the R10LB i-modulon activities were near the reference condition. The FliA i-modulon, which controls chemotaxis, was activated in all three media, whereas the FlhDC i-modulon, which controls biosynthesis and assembly of flagella, was strongly activated in RPMI (Figure S2d).

### Consistent transcriptional effects of subinhibitory antibiotic treatment

Since the ICA-based decomposition of transcriptomes provided a clear explanation of the complex responses to different media compositions, we used a similar approach to investigate the effects of antibiotic treatment at the subinhibitory concentrations presented in Figure 1.

The i-modulon activities for the antibiotic-treated conditions were primarily clustered by media, even when controlling for the media-specific responses (Figure 4a). Meropenem treatment on R10LB had minimal effects on the transcriptome and was removed from subsequent analyses.

**Figure 4:**
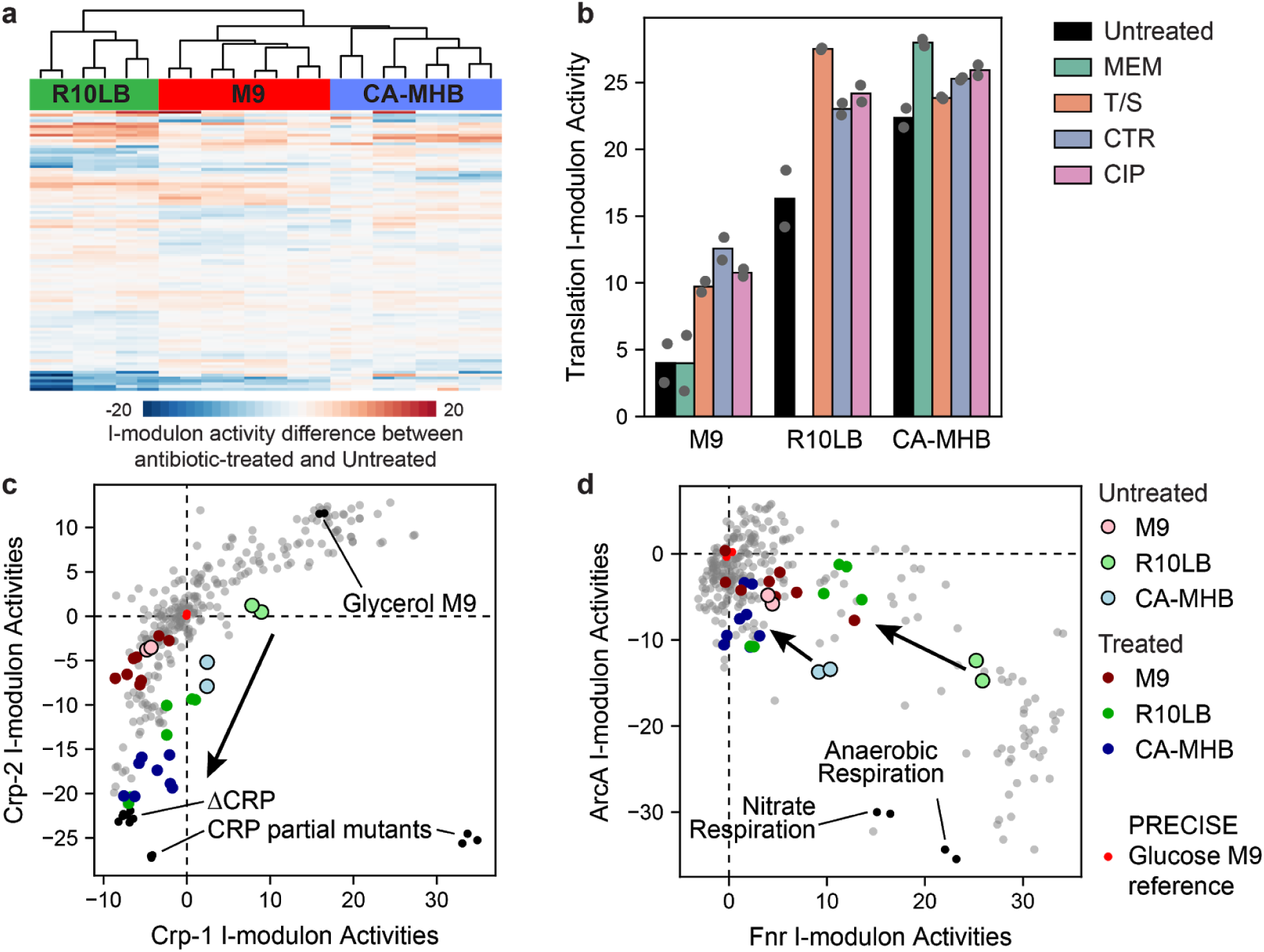
Consistent i-modulon responses to antibiotic treatment. (a) Clustered heatmap of expression differences in antibiotic-treated samples. Expression differences shown are between the antibiotic treated sample and the untreated sample in the same media condition. (b) Bar chart of translation i-modulon activities with and without antibiotic treatment. (c) Scatter plot of the Crp-1 and Crp-2 i-modulon activities across PRECISE. Arrow indicates the effect of antibiotic treatment in rich media. I-modulon activities from CRP knock-out and partial knock-out strains are also shown. (d) Scatter plot of Fnr and ArcA i-modulon activities across PRECISE. Arrows indicate the effect of antibiotic treatment in rich media. Anaerobic respiration conditions from PRECISE are also highlighted.

We first investigated common trends in i-modulon activities upon treatment by any antibiotic. Antibiotic treatment increased activity of the translation i-modulon in all media compositions (Figure 4b), but left the other Fear-Greed trade-off i-modulons largely unchanged (Figure S3).

Although none of the selected antibiotics directly interact with the ribosome, this indicated that antibiotic-mediated stress could indirectly increase ribosome levels in the cell.

We also observed significant reduction in both Crp-related i-modulon activities upon antibiotic treatment in rich media (Figure 4c). The Crp i-modulons encode genes in secondary carbon source catabolism; the Crp-1 i-modulon contains scavenging enzymes such as tryptophanase and acetyl-coA synthetase, whereas the Crp-2 i-modulon represents expression of phosphotransferase systems for alternative carbon sources.

Nutrient usage in the two rich media was nearly identical in the untreated samples, except for i-modulons encoding histidine biosynthesis (His-tRNA), branched-chain amino acid biosynthesis (Leu/Ile), and nucleotide biosynthesis (PurR-1 and PurR-2). However, upon treatment by any antibiotic, the activities of these i-modulons converged to similar levels in rich media (Figure S4).

### Subinhibitory antibiotic treatment decelerates cellular respiration in rich media

Across PRECISE, we observed a strong significant negative correlation between Fnr and ArcA i-modulon activities (Pearson R = −0.74, p-value < 10^−10^, Figure 4d), which represent two major respiration-related transcription factors. Fnr contains an iron-sulfur cluster that is destabilized in the presence of molecular oxygen (Kiley and Beinert, 1998), rendering Fnr inactive. Therefore, the high expression of these genes in R10LB indicates a high respiration rate that leaves little intracellular molecular oxygen available to destabilize Fnr’s iron-sulfur cluster. Herein lies one of the major differences between the two media; R10LB appears to promote a higher respiration rate than CA-MHB.

In both rich media, antibiotic treatment resulted in significant drops in Fnr i-modulon activities, indicating higher intracellular oxygen levels. This drop was coupled by a simultaneous increase in ArcA-regulated genes in the TCA cycle. Together, these two i-modulons point towards deceleration of the high respiratory rate facilitated by the rich media and a reduction in the redox state, as sensed by the ArcAB response system (Georgellis et al., 2001). A prior study used biochemical assays to investigate the effects of bacteriostatic antibiotics on *E. coli* (Lobritz et al., 2015), and observed that antibiotic treatment drastically reduced the respiration rate and moderately reduced the redox state of the cell. However, this effect seemed to be mitigated in minimal media, where we inferred a lower cellular respiration rate.

### Ciprofloxacin treatment leads to divergence in iron regulation

For most i-modulons, treatment by any antibiotic in subinhibitory concentrations resulted in a convergence of i-modulon activity between R10LB and CA-MHB. However, these mirrored responses do not explain why drugs like ciprofloxacin or meropenem have diverging MICs on the two media. Here, we will focus on ciprofloxacin and suggest a hypothesis towards its differential activity on the two media.

Ciprofloxacin blocks DNA gyrase, creating double-stranded DNA breaks (Chen et al., 1996). Subsequently, we observed a strong activity increase in the i-modulon representing the SOS-response regulator LexA in all three media (Figure 5a). In addition, ciprofloxacin has diverging effects on the two Fur-related i-modulons. The two Fur i-modulons contain many of the same genes, and exhibit a general trend in their activities across the entire PRECISE database (Figure 5b). The Fur-1 i-modulon responds to iron starvation, de-repressing iron (II) and (III) siderophore synthesis and transport systems, ribonucleotide reductases, superoxide dismutase, and iron-sulfur cluster assembly. The Fur-2 i-modulon responds to excess iron, further repressing siderophore transport and repressing the energy-transducing Ton system.

**Figure 5:**
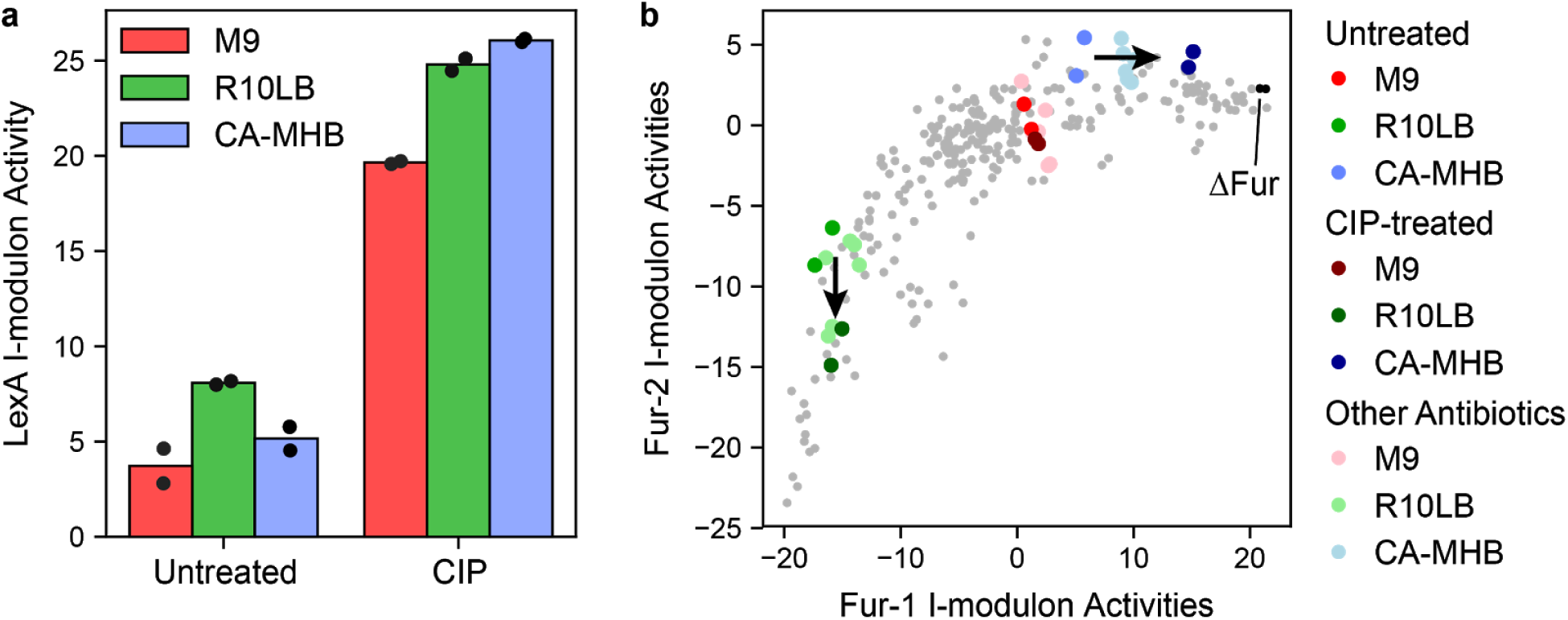
I-modulon responses to ciprofloxacin treatment. (a) The DNA-damage response regulator LexA is activated by ciprofloxacin treatment. (b) Scatter plot of the Fur-1 and Fur-2 i-modulon activities. Expression profiles from PRECISE are shown in light gray, and other colors are described in the legend. Arrows indicate i-modulon shifts in R10LB and CA-MHB resulting from ciprofloxacin treatment.

The Fur i-modulon activities diverge further upon ciprofloxacin treatment (Figure 5b), indicating an increase in free iron in R10LB and a stronger iron starvation response in CA-MHB. Between the increased intracellular oxygen levels observed in the previous section, and the increase in free iron specific to R10LB, the increased susceptibility of *E. coli* to ciprofloxacin in R10LB may be explained by Fenton chemistry, in which unincorporated iron and hydrogen peroxide generate hydroxyl radicals to further damage DNA (Imlay, 2013).

Prior studies have observed that reactive oxidative species, such as those created by Fenton chemistry, may contribute to the antibacterial action of ciprofloxacin (Dwyer et al., 2014, 2007; Goswami et al., 2006). However, it is not clear if iron levels directly affect ciprofloxacin efficacy, especially in light of the controversy regarding the role of oxidative stress in antibiotic lethality (Dwyer et al., 2015; Keren et al., 2013; Liu and Imlay, 2013). Additional studies investigating this relationship are therefore warranted.

## Discussion

Bacterial responses to antibiotics are complex, and depend on a variety of environmental and genetic factors. In this study, we explored the effects of different media compositions and antibiotic treatments on the transcriptomic state of the model organism *E. coli*. First, we showed that i-modulons could simplify the interpretation of large changes in expression due to varying environmental conditions. We then leveraged i-modulon activities with biochemical knowledge to infer the respiratory rate of *E. coli*, and subsequently elucidated a drop in respiration caused by antibiotic treatment. We also noted a significant difference in Fur activity between the two rich media, providing a potential hypothesis behind the differential susceptibility to ciprofloxacin in the two media.

We have presented an analytical pipeline for probing and contextualizing the transcriptomic effects of antibiotic treatment using a model organism. Although our analysis was focused on subinhibitory effects of antibiotics, this could be extended to investigate inhibitory or lethal doses of antibiotics. As publicly available transcriptomic datasets grow in size and diversity (Barrett et al., 2013), this pipeline can be applied to pathogenic strains and organisms grown in physiologically relevant media conditions. Future studies could explore the metabolic and transcriptional shifts associated with antibiotic treatment, leading to a “white-box” approach that connects antibiotic-treated transcriptomes to biochemical knowledge (Yang et al., 2019).

## Methods

### Bacterial strains and growth conditions

Escherichia coli strain MG1655 was grown in three media: (1) M9 minimal medium (47.8 mM Na2HPO4, 22 mM KH2PO4, 8.6 mM NaCl, 18.7 mM NH4Cl, 2 mM MgSO4 and 0.1 mM CaCl2) supplemented with 0.2% (w/v) glucose (M9); (2) Roswell Park Memorial Institute 1640 (RPMI) (Thermo Fisher Scientific) supplemented with 10% LB (R10LB); (3) Mueller Hinton Broth (Sigma-Aldrich) supplemented with 25 mg/L Ca^2+^ and 12.5 mg/L Mg^2+^ (CA-MHB). Glycerol stocks of E. coli were inoculated into the respective media and cultured overnight at 37 °C with vigorous agitation.

### Antibiotic Susceptibility Testing

Antibiotic susceptibility was determined as described previously (Dillon et al., 2019). Briefly, E. coli were cultured in the same media throughout (M9, R10LB, or CA-MHB) prior to the addition of antibiotics. Mid-logarithmic phase cultures were diluted to approximately ~2.5 × 10^7^ CFU/mL (~OD^600^ = 0.08) and diluted 1:100 in 200 μL media prepared with serial dilution. Plates were incubated with shaking at 200 rpm at 37°C overnight. Bacterial growth (OD600) was determined approximately 20 h later utilizing a Enspire Alpha multimode plate reader (PerkinElmer). To calculate the MIC90, defined as the amount of drug required to inhibit ≥90% of the growth of the untreated controls, the density of each drug-treated well was compared to untreated control. MIC for high-density cultures were performed as described above, except with a starting OD600~0.05 after dilution.

### Killing Assays

Mid-log *E. coli* MG1655 was used to inoculate 15 ml of either glucose M9, R10LB, or CA-MHB with approximately 2.5 ∗ 10^7^ CFU/mL (OD600~0.05). Each experimental well of the 96-well flat bottom plate received 180 μl of bacterial culture and 20 μl of the desired 10x drug stock. Plates were incubated shaking at 100 rpm at 37C overnight. After 20 h, plates were removed from the incubator and serial 10-fold dilutions of each well performed in their respective media. Twenty microliters of each serial dilution was spot plated onto LA and incubated at 37C overnight to enumerate the CFU.

### RNA-seq expression profiling and processing

Total RNA was sampled from duplicate cultures. Strains were cultured overnight in the respective media as described above. Mid-log phase cultures were diluted (starting OD600~0.05) into 30mL media for untreated conditions, or 30 mL media containing the MIC of the respective antibiotic for the respective media (Table 1). Flasks were incubated for 30 min at 37C with shaking. Cell broth was centrifuged and supernatant was removed. RNA extraction and library preparation were performed as described in (Sastry et al., 2019). Raw sequencing reads were performed as described in (Sastry et al., 2019). Differential expression was performed using DESeq2 (Love et al., 2014), with a log_2_ fold change cutoff of 1.5 and q-value cutoff of 0.05.

### ICA

Log-transformed transcripts per million (log-TPM) expression levels were concatenated to the PRECISE dataset. ICA and i-modulon processing were performed as described in (Sastry et al., 2019). Briefly, we executed FastICA 100 times with random seeds and a convergence tolerance of 10^−6^ for RNA-seq data, and a convergence tolerance of 10^−7^ for proteomics data. We constrained the number of independent components (ICs) in each iteration to the number of components that reconstruct 99% of the variance as calculated by principal component analysis. The resulting ICs were clustered using DBSCAN to identify robust ICs, with parameters with epsilon of 0.1, and minimum cluster seed size of 50. This process was repeated 10 times, and only ICs that consistently occurred in all runs were kept.

As described in Sastry et al.(Sastry et al., 2019), i-modulons were extracted from ICs by iteratively removing genes with the largest absolute value and computing the D’agostino K2 test statistic of the resulting distribution. Once the test statistic fell below a cutoff, which was identified through a sensitivity analysis (Sastry et al., 2019), we designated the removed genes as the “i-modulon”.

### Explained variance

Explained variance between two conditions was calculated as follows:

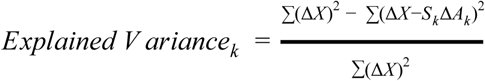

Where *k* is the i-modulon of interest. The total explained variance was calculated similarly:

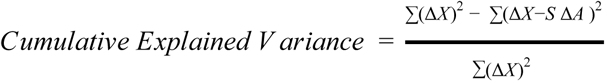

The difference between the CEV and the sum of the explained variance is the confounding variance.

## Acknowledgements

The authors would like to thank George Sakoulas, Hannah Tsunemoto, Yara Seif, and CJ Norsigian for informative discussions. This research used resources of the National Energy Research Scientific Computing Center, a DOE Office of Science User Facility supported by the Office of Science of the U.S. Department of Energy under Contract No. DE-AC02-05CH11231. This work was funded by the Novo Nordisk Foundation Center for Biosustainability and the Technical University of Denmark (grant number NNF10CC1016517) and by the NIH NIAID (grant number 1-U01-AI124316).

## Notes

### Competing Interest Statement

The authors have declared no competing interest.

## References

Anand A, Chen K, Catoiu E, Sastry AV, Olson CA, Sandberg TE, Seif Y, Xu S, Szubin R, Yang L, Feist AM, Palsson BO. 2019. OxyR is a convergent target for mutations acquired during adaptation to oxidative stress-prone metabolic states. Mol Biol Evol. doi:10.1093/molbev/msz251

Barrett T, Wilhite SE, Ledoux P, Evangelista C, Kim IF, Tomashevsky M, Marshall KA, Phillippy KH, Sherman PM, Holko M, Yefanov A, Lee H, Zhang N, Robertson CL, Serova N, Davis S, Soboleva A. 2013. NCBI GEO: archive for functional genomics data sets—update. Nucleic Acids Res 41:D991–D995.

Chen CR, Malik M, Snyder M, Drlica K. 1996. DNA gyrase and topoisomerase IV on the bacterial chromosome: quinolone-induced DNA cleavage. J Mol Biol 258:627–637.

Dillon N, Holland M, Tsunemoto H, Hancock B, Cornax I, Pogliano J, Sakoulas G, Nizet V. 2019. Surprising synergy of dual translation inhibition vs. Acinetobacter baumannii and other multidrug-resistant bacterial pathogens. EBioMedicine 46:193–201.

Dorschner RA, Lopez-Garcia B, Peschel A, Kraus D, Morikawa K, Nizet V, Gallo RL. 2006. The mammalian ionic environment dictates microbial susceptibility to antimicrobial defense peptides. The FASEB Journal 20:35–42.

Dwyer DJ, Belenky PA, Yang JH, MacDonald IC, Martell JD, Takahashi N, Chan CTY, Lobritz MA, Braff D, Schwarz EG, Ye JD, Pati M, Vercruysse M, Ralifo PS, Allison KR, Khalil AS, Ting AY, Walker GC, Collins JJ. 2014. Antibiotics induce redox-related physiological alterations as part of their lethality. Proc Natl Acad Sci U S A 111:E2100–9.

Dwyer DJ, Collins JJ, Walker GC. 2015. Unraveling the physiological complexities of antibiotic lethality. Annu Rev Pharmacol Toxicol 55:313–332.

Dwyer DJ, Kohanski MA, Hayete B, Collins JJ. 2007. Gyrase inhibitors induce an oxidative damage cellular death pathway in Escherichia coli. Mol Syst Biol 3:91.

Ersoy SC, Heithoff DM, Barnes L, Tripp GK, House JK, Marth JD, Smith JW, Mahan MJ. 2017. Correcting a Fundamental Flaw in the Paradigm for Antimicrobial Susceptibility Testing. EBioMedicine 20:173–181.

Farha MA, French S, Stokes JM, Brown ED. 2018. Bicarbonate Alters Bacterial Susceptibility to Antibiotics by Targeting the Proton Motive Force. ACS Infect Dis 4:382–390.

Federowicz S, Kim D, Ebrahim A, Lerman J, Nagarajan H, Cho B-K, Zengler K, Palsson B. 2014. Determining the control circuitry of redox metabolism at the genome-scale. PLoS Genet 10:e1004264.

Georgellis D, Kwon O, Lin EC. 2001. Quinones as the redox signal for the arc two-component system of bacteria. Science 292:2314–2316.

Goswami M, Mangoli SH, Jawali N. 2006. Involvement of reactive oxygen species in the action of ciprofloxacin against Escherichia coli. Antimicrob Agents Chemother 50:949–954.

Imlay JA. 2013. The molecular mechanisms and physiological consequences of oxidative stress: lessons from a model bacterium. Nat Rev Microbiol 11:443–454.

Keren I, Wu Y, Inocencio J, Mulcahy LR, Lewis K. 2013. Killing by Bactericidal Antibiotics Does Not Depend on Reactive Oxygen Species. Science. doi:10.1126/science.1232688

Kiley PJ, Beinert H. 1998. Oxygen sensing by the global regulator, FNR: the role of the iron-sulfur cluster. FEMS Microbiol Rev 22:341–352.

Kolb A, Busby S, Buc H, Garges S, Adhya S. 1993. Transcriptional regulation by cAMP and its receptor protein. Annu Rev Biochem 62:749–795.

Kudrin P, Varik V, Oliveira SRA, Beljantseva J, Del Peso Santos T, Dzhygyr I, Rejman D, Cava F, Tenson T, Hauryliuk V. 2017. Subinhibitory Concentrations of Bacteriostatic Antibiotics Induce relA-Dependent and relA-Independent Tolerance to β-Lactams. Antimicrob Agents Chemother 61. doi:10.1128/AAC.02173-16

Lin L, Nonejuie P, Munguia J, Hollands A, Olson J, Dam Q, Kumaraswamy M, Rivera H, Corriden R, Rohde M, Hensler ME, Burkart MD, Pogliano J, Sakoulas G, Nizet V. 2015. Azithromycin Synergizes with Cationic Antimicrobial Peptides to Exert Bactericidal and Therapeutic Activity Against Highly Multidrug-Resistant Gram-Negative Bacterial Pathogens. EBioMedicine. doi:10.1016/j.ebiom.2015.05.021

Liu Y, Imlay JA. 2013. Cell death from antibiotics without the involvement of reactive oxygen species. Science 339:1210–1213.

Lobritz MA, Belenky P, Porter CBM, Gutierrez A, Yang JH, Schwarz EG, Dwyer DJ, Khalil AS, Collins JJ. 2015. Antibiotic efficacy is linked to bacterial cellular respiration. Proc Natl Acad Sci U S A 112:8173–8180.

Love MI, Huber W, Anders S. 2014. Moderated estimation of fold change and dispersion for RNA-seq data with DESeq2. Genome Biol 15:550.

Meylan S, Andrews IW, Collins JJ. 2018. Targeting Antibiotic Tolerance, Pathogen by Pathogen. Cell 172:1228–1238.

Poudel S, Tsunemoto H, Seif Y, Sastry A, Szubin R, Xu S, Machado H, Olson C, Anand A, Pogliano J, Nizet V, Palsson BO. 2020. Revealing 29 sets of independently modulated genes in Staphylococcus aureus, their regulators and role in key physiological responses. bioRxiv. doi:10.1101/2020.03.18.997296

Rychel K, Sastry AV, Palsson B. 2020. Machine learning uncovers independently regulated modules in the Bacillus subtilis transcriptome. bioRxiv. doi:10.1101/2020.04.26.062638

Saelens W, Cannoodt R, Saeys Y. 2018. A comprehensive evaluation of module detection methods for gene expression data. Nat Commun 9:1090.

Sanchez-Vazquez P, Dewey CN, Kitten N, Ross W, Gourse RL. 2019. Genome-wide effects on Escherichia coli transcription from ppGpp binding to its two sites on RNA polymerase. Proc Natl Acad Sci U S A 116:8310–8319.

Sastry AV, Gao Y, Szubin R, Hefner Y, Xu S, Kim D, Choudhary KS, Yang L, King ZA, Palsson BO. 2019. The Escherichia coli transcriptome mostly consists of independently regulated modules. Nat Commun 10:5536.

Seo SW, Kim D, Latif H, O’Brien EJ, Szubin R, Palsson BO. 2014. Deciphering Fur transcriptional regulatory network highlights its complex role beyond iron metabolism in Escherichia coli. Nat Commun 5:4910.

Seo SW, Kim D, O’Brien EJ, Szubin R, Palsson BO. 2015. Decoding genome-wide GadEWX-transcriptional regulatory networks reveals multifaceted cellular responses to acid stress in Escherichia coli. Nat Commun 6:7970.

Utrilla J, O’Brien EJ, Chen K, McCloskey D, Cheung J, Wang H, Armenta-Medina D, Feist AM, Palsson BO. 2016. Global Rebalancing of Cellular Resources by Pleiotropic Point Mutations Illustrates a Multi-scale Mechanism of Adaptive Evolution. Cell Syst 2:260–271.

Yang JH, Wright SN, Hamblin M, McCloskey D, Alcantar MA, Schrübbers L, Lopatkin AJ, Satish S, Nili A, Palsson BO, Walker GC, Collins JJ. 2019. A White-Box Machine Learning Approach for Revealing Antibiotic Mechanisms of Action. Cell 177:1649–1661.e9.

Zampieri M, Zimmermann M, Claassen M, Sauer U. 2017. Nontargeted Metabolomics Reveals the Multilevel Response to Antibiotic Perturbations. Cell Rep 19:1214–1228.

