## Supplementary Figures for "Decomposition of transcriptional responses provides insights into differential antibiotic susceptibility"

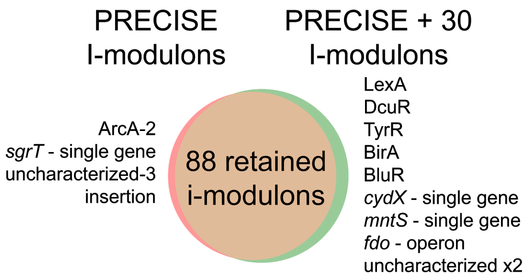


**Figure S1:** Venn diagram comparing i-modulons generated from PRECISE compared to i-modulons generated from PRECISE + 30 new expression profiles

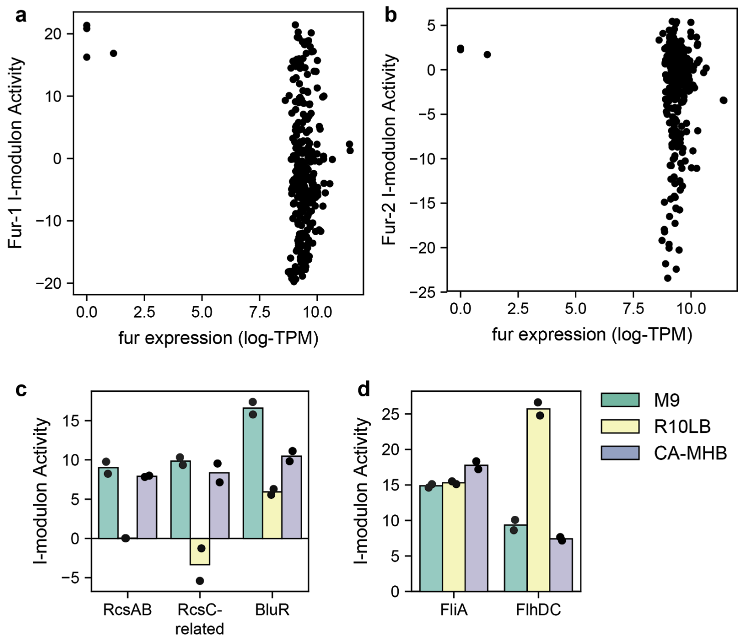


**Figure S2:** Additional information relating to Figure 3. (a,b) Scatterplot between the Fur i-modulons and iron concentrations. (c,d) I-modulon activities of rcs-phosphorelay-related i-modulons and motility i-modulons across three media conditions.

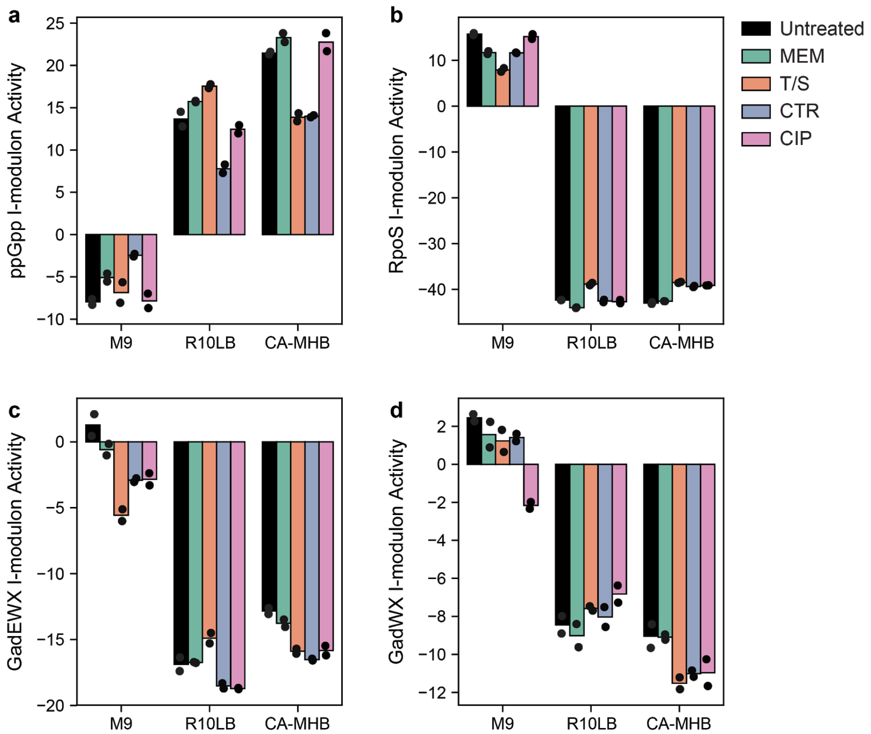


**Figure S3**: Bar chart of fear-greed i-modulon activities, including the (a) ppGpp i-modulon; (b) RpoS i-modulon; (c) GadEWX i-modulon; (d) GadWX i-modulon.

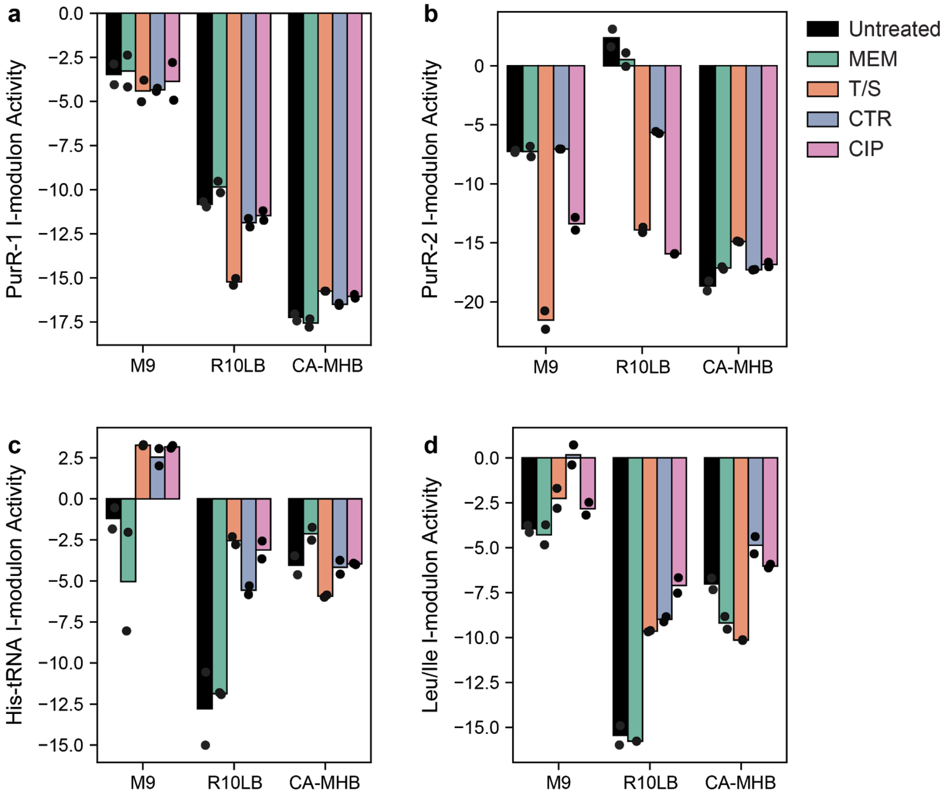


**Figure S4**: Bar chart of nutrient-related i-modulon activities, including the (a) PurR-1 i- modulon; (b) PurR-2 i-modulon; (c) His-tRNA i-modulon; (d) Leu/Ile i-modulon.
